# Accurate Filtering of Privacy-Sensitive Information in Raw Genomic Data

**DOI:** 10.1101/292185

**Authors:** Jérémie Decouchant, Maria Fernandes, Marcus Völp, Francisco M Couto, Paulo Esteves-Veríssimo

## Abstract

Sequencing thousands of human genomes has enabled breakthroughs in many areas, among them precision medicine, the study of rare diseases, and forensics. However, mass collection of such sensitive data entails enormous risks if not protected to the highest standards. In this article, we follow the position and argue that post-alignment privacy is not enough and that data should be automatically protected as early as possible in the genomics workflow, ideally immediately after the data is produced. We show that a previous approach for filtering short reads cannot extend to long reads and present a novel filtering approach that classifies raw genomic data (i.e., whose location and content is not yet determined) into privacy-sensitive (i.e., more affected by a successful privacy attack) and non-privacy-sensitive information. Such a classification allows the fine-grained and automated adjustment of protective measures to mitigate the possible consequences of exposure, in particular when relying on public clouds. We present the first filter that can be indistinctly applied to reads of any length, i.e., making it usable with any recent or future sequencing technologies. The filter is accurate, in the sense that it detects all known sensitive nucleotides except those located in highly variable regions (less than 10 nucleotides remain undetected per genome instead of 100,000 in previous works). It has far less false positives than previously known methods (10% instead of 60%) and can detect sensitive nucleotides despite sequencing errors (86% detected instead of 56% with 2% of mutations). Finally, practical experiments demonstrate high performance, both in terms of throughput and memory consumption.

## I. BACKGROUND

In recent years, next-generation sequencing technologies have evolved to produce biological data faster and with improved quality. Those technologies are now well developed for short reads (DNA sequences of less than 100 nucleotides), however, a new generation producing longer reads has already entered the market [1]. In practice, those reads are often aligned to a reference genome to obtain their location in that reference, and then infer genomic insights (e.g., variant calling, disease development risk, etc.) [2], [3].

Human DNA sequences have a high value to biomedical research if studied at a large scale, as they increase the accuracy of studies and enable discoveries of new variants. However, the number of human DNA sequences available for research is still limited. Some outstanding initiatives, such as the 1000 genomes project, provide open access to human genomes, but the number of accessible genomes is quite limited when compared to the number of genomes being sequenced daily. In addition, researchers typically have personalized access to specific biobanks, which is granted under the terms-of-use defined by these banks. However, biobanks are often limited to a specific population, hindering large-scale diversity studies. Widely sharing genomes would also improve the reproducibility of results, since studies based on information from biobanks cannot be reproduced easily due to biobanks tightly controlling access.

Risks to privacy are still an obstacle to sharing. Several attacks against genomic privacy have been described in the literature. Using genomic information, such as single nucleotide polymorphisms (SNPs), short tandem repeats, disease related genes, and possibly different kinds of publicly available personal details, these attacks fit into one of several categories: re-identification attacks [4], [5], [6], [7], [8], recovery attacks [9], and membership attacks [10], [11], [12]. These genomic privacy attacks alerted the research community for the need to replace the conventional procedures by privacy-preserving frameworks. In the last years, several protocols have been designed to protect the sensitive information contained in genomic data [13], [14], [15], [4], [16]. Concerning the read alignment step, secure cryptographic algorithms have been designed to align reads while protecting their content [17], [18]. However, their slow performance fails to match the throughput of current sequencing machines at manageable cost, which leads biological research centers to continue to rely on plaintext alignment.

Future genomics research would benefit from software solutions capable of protecting sensitive information throughout its whole life-cycle. That is, from its initial acquisition during the sequencing phase, to the day it is not sensitive anymore, including its storage and use by researchers or physicians. Secure alignment algorithms are promising tools that could be integrated in such a secure environment, which would therefore protect all raw genomic information preventively. Then, after the alignment step sensitive information would be identified, since aligned reads reveal the role of sequences in the DNA, and the protection level of non-sensitive data adapted. However, applying such a paradigm implies a high performance overhead since secure alignment algorithms are slower than the plaintext ones.

This manuscript proposes an alternate method, which detects genomic variations in the early stages of the genomic pipelines. This approach is based on the existing databases of genomic variations, which are getting more and more stable [15], and makes use of Bloom filters for space efficiency and fast membership testing. Indeed, as previously argued and as we independently confirmed, the rate of new discoveries has been decreasing over the years, which makes our approach very timely. For example, the number of new SNPs discovered in the 1000 genomes project has been divided by 120 between 2010 (12 millions) and 2016 (100 thousands). We believe that such decrease is bound to be confirmed in the future.

We believe our approach to be an important step towards enforcing privacy prior to the alignment of reads, which is routinely executed in a public cloud for performance and cost reasons. Typically, clouds do not guarantee that the data they host cannot be accessed by either the cloud service provider, or an intruder [19], and several works highlighted the risks associated to the inconsiderate use of clouds for biomedical data [20], [21], [22], [23]. We therefore consider an adversary located in the cloud whose goal is to perform a privacy attack using raw genomic sequences it is able to observe either prior, or during, alignment.

We developed a filtering approach, which partitions raw genomic data into privacy sensitive or non-sensitive genomic data, and that is envisioned to enable the accurate, lightweight, and fast obfuscation of reads so that they can safely be manipulated in plaintext in clouds. The filter is lightweight, and fast enough, that it does not have to be parallelized in massive resources. The non-sensitive genomic data (i.e., obfuscated reads, which by definition are identical between any two subjects) are allowed to leave the secure environment (e.g., the sequencing machine) to be aligned in a public cloud, while the sensitive genomic data stays protected, in a private environment. In addition, we believe that an alignment scheme that would manipulate obfuscated versions of reads is feasible, even though, the precise design and evaluation of this scheme is future work. Relying on the filter to implement such an alignment scheme would result in a performance improvement compared to current cryptographic alignment schemes, since plaintext alignment is much faster. This manuscript focuses on the keystone of this envisioned scheme, a filter which accurately detects sensitive information inside raw genomic data, while future work will optimize the alignment of masked reads.

To the best of our knowledge, the only previous early filtering method [15] is limited to short reads (30 nucleotides) and to classifying them as privacy sensitive, or non-sensitive, depending on whether or not they are susceptible to carry a relevant genomic variation. As we explain in A, the mechanisms described in this previous work cannot be successfully used with long reads as they would produce too many false positives, since the proportion of reads that contain at least one sensitive nucleotide grows very quickly with their length. For example, 88% of the 100-nucleotide reads and 94% of the 200-nucleotide reads would be considered sensitive by this filtering method, which neutralizes its effectiveness.

This work is the result of an analysis of possible Bloom filter-based approaches to identify sensitive nucleotides from reads. Our analysis lead to the design of novel algorithms to filter privacy sensitive information from raw genomic data. We show through comprehensive experiments that the overall process improves upon the state of the art as follows:

1. it can be applied on reads of any size, output by past, and current sequencing technologies, which produce longer and for now, more error-prone reads;
2. it is designed to detect all kinds of sensitive genomic variations (i.e., including insertions/deletions) and miss less than 10 sensitive nucleotides per genome (instead of several thousands), yielding high privacy protection;
3. it considerably improves on previous filtering false positive rate at the nucleotide level (from 60% in previous works to 10%);
4. multiple filters can be used in parallel to tolerate more sequencing errors (86% of the sensitive nucleotides are still detected with 2% sequencing errors; previous works detect 56% in this situation). Error tolerance eases an early adoption while sequencing technologies continue to improve their detection accuracy.

### A. Related work

#### 1) Privacy attacks

Recently, numerous privacy attacks have demonstrated the sensitivity of genomic, and more generally, biological data. Homer et al. showed that even a very small proportion in aggregate allele frequencies dataset is sufficient to identify contributing individuals using their DNA profile [10]. This attack was later refined by Wang et al. in [4]. In particular, they showed that only 100 SNPs can be enough to reidentify a subject. It has also been shown that 92 well-chosen SNPs are enough to identify anyone in the world with high probability [5]. Kidd et al. [6] built a panel of 19 SNPs that, with high probability, can be used to accurately determine the ethnic origin of subjects. Goodrich showed that successive queries over encrypted data may allow an attacker to gain access to private data [24]. In [11], Shringarpure and Bustamante showed that in a beacon made of 65 subjects from the 1000 genomes project, querying 250 SNPs is enough to re-identify a subject. Gymreck et al. re-identified the individuals behind 131 genomes contributing their DNA to the 1000 genome project [7]. More recent attacks studied the sensitivity of other types of biological data. For example, it has been shown that miRNA data can be used in membership attacks [12].

#### 2) Genomic privacy preserving mechanisms

Genomic privacy is considered one of the major challenges of the biomedical community [25], [26], [27], [28].There are three different ways to protect the privacy of genomic data: (i) data deidentification [29], [30]; (ii) data augmentation [31], [32]; and (iii) methods based on cryptographic methods [24], [33].

The data deidentification methods tend to remove or encrypt personal identifiers, such as social security numbers, zip codes, or names, which are initially associated with genomic data. Nevertheless, these methods cannot guarantee sufficient privacy protection and are not able to deal with reidentification problems [10], [34], [7], [11].

Data augmentation methods achieve the goal of privacy protection by generalizing, each record so that it cannot be distinguished from some other shared records [31], [35], [32]. With this kind of methods, the privacy of genomic data can be enforced, at the expense of controlled loss of utility.

Cryptology-based methods do not access the original data. They maintain data utility by using privacy-preserving data querying methods that can be applied to genomic sequences [24], [33]. Protection methods based solely on cryptographic algorithms are not sufficient, since encryption mechanisms can be broken in a comparably shorter time than the personal genomic privacy protection requires [36].

The work we describe in this manuscript can be classified into the data augmentation methods, since our filter aims at detecting and (temporarily) obfuscating sensitive nucleotides from DNA sequences prior to the read alignment step.

#### 3) Early read filtering

The work we present is this manuscript focus on read filtering, i.e., the early detection of sensitive nucleotides in reads at the mouth of, and possibly inside, the sequencing machines. Requirements for an early read filtering method include i) to be accurate (i.e., correctly detect sensitive nucleotides); ii) not to be a throughput bottleneck (i.e., it must filter reads faster than the sequencing machine produces them); and iii) to be practical (i.e., it can be implemented in, or close to, a sequencing machine, with limited hardware resources).

To the best of our knowledge, the only previous early read filtering approach, the short read filtering (SRF) approach, has been presented in [15]. The SRF describes the filtering of privacy sensitive information contained in short sequences (30 nucleotides) of raw genomic data. SRF uses a Bloom filter [37], which is a data structure that is used to test whether a given element is part of a predefined set, to determine if a short read is part of a previously built dictionary of known sensitive reads. Therefore, SRF first creates a dictionary of all sensitive sequences made of either 30 nucleotides. In practice, a Bloom filter is implemented using a bit table and a set of hash functions. To initialize the bit table, each sequence in the dictionary is hashed, using the hash functions, to produces bit indexes, and set the corresponding bits to 1. Testing whether a read is part of the bit table then consists in checking that all bits that the hash functions map it to are set to 1. Thanks to its Bloom filter, SRF produces few false positives, always reidentifies known sensitive sequences, and maintains a high throughput.

However, as we explain in A, until now, the filtering of long reads was an open challenge, as SRF extended in the most efficient way to long reads would consider 88% of the 100-nucleotides reads and 94% of the 200-nucleotides sensitive, rendering it useless for any possible application.

#### 4) Secure read alignment

Aligning reads in an unsecure environment, such as a public cloud, has clearly been identified as a threat [38]. More precisely if the adversary is able to align those observed reads and identify the genomic variations, a trail attack becomes a possibility [30]. In a trail attack, DNA samples are matched to their identified subject through the use of unique distinguishing features in different institutions where the subject leaves traces of his genome. For example, if a subject, whose reads have been aligned in an attacked cloud, later participates in a public biomedical study, the adversary may be able to identify the subject in the results based on the subjects’ known genomic variations, and infer private information (e.g., a disease associated with the study, or the subject’s identity). Read alignment is therefore the first stage in the genomic pipeline where privacy-sensitive genomic features can be observed by an adversary, if they are not protected.

Secure read alignment is related to read filtering, but is a different problem. Secure alignment algorithms aim at aligning reads to a reference genome while preventing an adversary to observe the sensitive nucleotides contained in reads. Read filtering would reinforce the reads security during their storage prior to their alignment. In addition, novel read alignment schemes may potentially be designed by building on read filtering methods, however, this is future work.

Among the existing solutions presented, Zhang et al. [39] described a hybrid cloud-based solution, which requires a private/non-private data classifier to benefit from the advantages of a public and a private clouds, while limiting their respective drawbacks. The long read filter could be that classifier. Closely related, Chen et al. [40] described a distributed approach that is based on hashing short substrings of reads (seeds), and mapping them with the reference, relying on both a public and a private cloud. This approach keeps the sensitive data processing in the private cloud, and the most exhaustive computations in the public cloud. At the time this manuscript was written, Popic et al. argued that this approach does not prevent some Kmers repeats to be observed, and proposed Balaur [41], a privacy-preserving alignment protocol based on locality-sensitive hashing. Balaur relies on encryptions and on a large amount of memory. Finally, secure alignment algorithms proven to be secure are either based on garbled-circuit [18], or on homomorphic encryption [16], [42]. However, their wide use is currently limited by their low throughput and their high communication cost.

#### 5) Read obfuscation

The early detection and protection of privacy sensitive information is a goal that the research community is actively trying to address [43], [44]. Previous works have argued for the partitioning of genomic data into a sensitive and a non-sensitive part, but assume it is either done manually [13] or using a hypothetical external tool [14]. The approach we describe in this paper is fully automatic, and works on the earliest form of genomic data.

In [13], Ayday et al. present a distributed architecture that allows medical units to query the DNA data of a patient from aligned reads, while avoiding to reveal sensitive SNPs. However, the process described in this paper is not automatic, since sensitive portions of the DNA have to be declared by scientists. This mechanism also concerns a later stage of the DNA workflow, where genomic reads have been aligned, and sensitive information exactly identified.

Several other works proposed computational methods for anonymizing databases of DNA sequences by obfuscating sensitive nucleotides. Malin et al. [32] presented a DNA sequence anonymity method called DNA lattice anonymization (DNALA), which enforces the k-anonymity [45] privacy property. Li et al. [46] further improved performance by replacing CLUSTALW [47], the multiple sequence alignment software used in DNALA, with MegaBLAST [48], a global pair-wise alignment software. Wan et al. [49] further iterates the obfuscation process with MegaBLAST until all sequences in the dataset are clustered and obfuscated, and claim that it improves the utility accuracy and performance.

Compared to the filter we describe in this manuscript, those approaches do not satisfy all the criteria we listed for early read filtering. First, they require consequent computational resources to identify differences between reads. In contrast, our filter is lightweight, and fast enough, as we show, that it does not have to be parallelized in massive resources, and can thus be used, for example, in a secluded data center integrated with the sequencing machine. In addition, these works aim at anonymizing a database of reads, which is a different goal than filtering a subject’s reads. Using these approaches to detect all sensitive nucleotides in a subject’s reads would require the alignment of those reads to all possible sensitive reads, which is clearly much more costly than our approach.

#### 6) Bloom filters

We refer to [50] for a survey on Bloom filters [37], and their various possible uses. Bloom filters are implemented using a bit table and hash functions that map an element to bit indices. The initialization of a Bloom filter with a set consists in computing the bit indexes for all elements in the set, and setting all corresponding bits to 1. Later, testing whether an element is part of the initialization set consists in checking that all bits that the hash functions map it to are set to 1.

A key characteristic of Bloom filters is their absence of false negatives. However, they may produce false positives (i.e., they may answer that an element is part of the initialization set when in reality it is not). The false positive rate *p* is usually fixed by the user, at Bloom filter parametric initialization time, along with the number of elements *n* that are to be inserted. The other implicit parameters of a Bloom filter, namely its size *m* (in bits) and the number of hash functions *k* it uses, can then be derived from those initialization values, and more specifically, using the two following asymptotic equations: 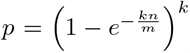 and 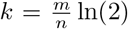. From these equations, one could derive that reducing a Bloom filter’s false positive rate, given an unmodified initialization set, implies to increase its size.

Bloom filters can be used either for space efficiency, or for privacy-preserving matching. For example, in the genomic privacy literature, they have been used for privacy preserving record linkage, which consist in linking a research subject’s information across different institutions. Existing works in that area include [51], [52], [53]. However, it may be unsecure to rely on Bloom filters for privacy, as some works highlighted [54] or addressed [55], [56].

In this manuscript, we rely on Bloom filters for space efficiency and high throughput reasons. More particularly, Bloom filters compactly represent data sets of sensitive sequences, and quickly answer membership queries (i.e., is a read known as being sensitive?). There are no privacy risks associated to our use of Bloom filters, since they are initialized with publicly known data.

## II. METHODS

In the following, we call sensitive either a nucleotide that is involved in a genomic variation or in a Short Tandem Repeat (STR), or a genomic sequence which contains at least one sensitive nucleotide. We call (*K, i*)-sensitive a sequence of *K* nucleotides where the *i*^*th*^ nucleotide is sensitive.

This section introduces the long read filtering approach (LRF). Section II-B explains how, given *K* and *i* values, we create a dictionary of every (*K, i*)-sensitive sequence that can be encountered in a human genome. Section II-C presents the Long Read Filtering (LRF) approach, which consists in studying all subsequences of a read using Bloom filters, initialized with dictionaries of (*K, i*)-sensitive sequences, to identify genomic variations. Finally, Section II-D extends this approach to tolerate sequencing errors (insertions, deletions or mutations) in reads.

### A. Data

We use the genomic variations and the individual genomes from the 1000 Genomes project [57], along with the short tandem repeats (STRs) from the Tandem Repeats Database [58] to create the dictionaries of sensitive sequences, and filter individual genomes. We use the Phase 3 20130502 release of the 1000 genomes project, and the associated GRCh38 reference project.

### B. Generating dictionaries of sensitive sequences

Generating all possible sensitive sequences, from either genomic variations or STRs, is a very important step for the accuracy of the filter. As previously said, the filter relies on dictionaries built from sequences that have a sensitive nucleotide (which can be part of genomic variation that involve more than one nucleotide) in a common position. We denote by *dict*_*i*_ the dictionary of (*K, i*)-sensitive sequences, i.e., sequences of *K* nucleotides whose *i*^*th*^ nucleotide is sensitive. In this section, we explain how we create these sequences from the 1000 Genomes project and the Tandem Repeats database.

#### 1) Genomic variations

Dictionaries of sensitive sequences are built depending on two parameters: the length *K* of the sequences, and a position 0 ≤ *i < K* at which all sequences will have a sensitive nucleotide (they may have other sensitive nucleotides). The generation of a dictionary can be summarized as follows. For every genomic variation in the 1000 Genomes Project, we first collect other genomic variations that are at a close distance from it, either located before or after in the human genome. Since individuals present combinations of several genomic variations, located at different locations, we then enumerate all theoretically possible combinations of the collected genomic variations. Finally, we create sequences from these combinations by relying on the reference genome to populate the gaps between the genomic variations. Finally, depending on the positions of the sensitive nucleotides in the sequences of the desired dictionary, i.e., depending on *i*, we then extract all adequate subsequences of length *K*.

When generating a sensitive sequence around a studied genomic variation, one has first to be careful to go far enough to collect genomic variations, since insertions/deletions may change the distances between sensitive nucleotides.

If it is possible, we then generate all combinations of the versions of the genomic variations. However, we observed that some sets of selected genomic variations contained up to 50 different locations, and since there are theoretically 2^*N*^ combinations of *N* genomic variations, it is not possible to generate all of them. In such cases, we selected the major allele for the genomic variations whose least frequent allele is the rarest among all remaining genomic variations until there remained 8 variations. We present results in A that justify why combining up to 8 variations from a genomic database allow us to detect all variations of a subject in practice.

Figure 1 illustrates the combination process for one genomic variation located at position *P*_2_ in the human genome, and for sequences of *K* = 8 nucleotides. In this example, two genomic variations are located less than *K* nucleotides away from position *P*_2_ when combined, namely at positions *P*_1_ and *P*_3_. Each of those genomic variations has two alleles: the reference one (represented in blue), and the alternative one (represented in red or green). From the reference genome, and the VCF file that contains the information about the genomic variations, our code generates all possible combinations of all three alleles, and then selects the adequate sensitive sequences depending on the dictionaries that need to be created. In this example, two dictionaries, *Dict*_0_ and *Dict*_29_, respectively contain the sensitive sequences whose first, respectively 30-th nucleotide, is involved in a genomic variation.

**Fig. 1.**
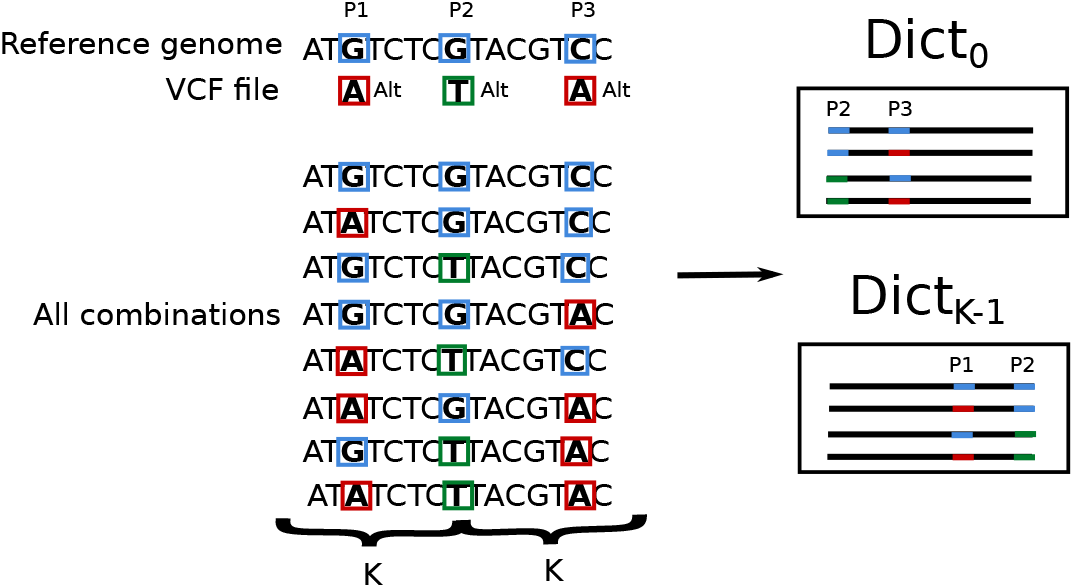
Illustration of the generation of dictionaries of sensitive sequences, of length K, from a genomic region that carries multiple SNPs.

#### 2) Short Tandem Repeats

Generating the sensitive sequences from STRs follow a similar pattern. Given the position of a STR, genomic variations located before and after it are collected, and combined. In the normal case, for all possible permutations of an STR pattern, the permutated pattern is concatenated until the concatenated sequence’s length is larger than *K* Then, the left and right flankings are generated from the reference genome, and from combining the genomic variations they contain. Finally, some sensitive sequences of *K* nucleotides are selected to be inserted in the adequate dictionaries.

However, it also happens that some STRs are never repeated sufficiently often for the repeated pattern to account for at least *K* nucleotides, which would prevent the previous method from generating the right subsequences. Consequently, for these short STR patterns with few repeats we start generating sensitive sequences with the minimal amount of repeats, and then increase the number of repeats progressively until either we reach the maximal number of repeats or the length of the repeated section accounts for more than *K* nucleotides. After each concatenation of a repeat pattern, we generate all possible (*K, i*)-sensitive sequences.

#### C. Long read filter

The long read filter creates one or several Bloom filters, from the previously generated dictionaries of (*K, i*)-sensitive sequences. Then, passing over the reads and filtering all sequences in a sliding window of size *K*, it checks whether each K-mer has been inserted in a Bloom filter, and therefore contains a sensitive character at the sensitive position of sequences of the corresponding dictionary (i.e., the *i* values of the (*K, i*)-sensitive sequences). This way, upon detection of a sequence in a Bloom filter only one nucleotide is detected sensitive. As we explain in A, previous and other Bloom filter based approaches would lead to much higher false positive rates than this one.

Figure 2 illustrates the LRF approach, where one thread, *Thread*_0_, detects sensitive nucleotides in a read, which has been given in input. This thread first initializes a Bloom filter *BF*_0_ with a dictionary, *Dict*_0_, of all possible 30-bases long sequences (i.e., here *K* = 30) whose first nucleotide is sensitive. The Bloom filter uses three hash functions, *h*_0_, *h*_1_ and *h*_2_. *Thread*_0_ is shown progressing in the read as it studies all 30-bases long subsequences of the read. The thread uses the same hash functions *h*_0_, *h*_1_ and *h*_2_, that were used to initialize *BF*_0_ to efficiently check if they are part of the dictionary *Dict*_0_. If that is the case, i.e., if all bits the hash functions lead to are set to 1, it tags the first nucleotide of the current subsequence as sensitive. In Figure 2, nucleotides detected sensitive are colored in red, and the output of the filter consists of the input read along with a set of the nucleotides flagged as sensitive. Revealing the non-sensitive nucleotides of a subject does not reveal any distinctive information, since by definition they are common to all subjects.

**Fig. 2.**
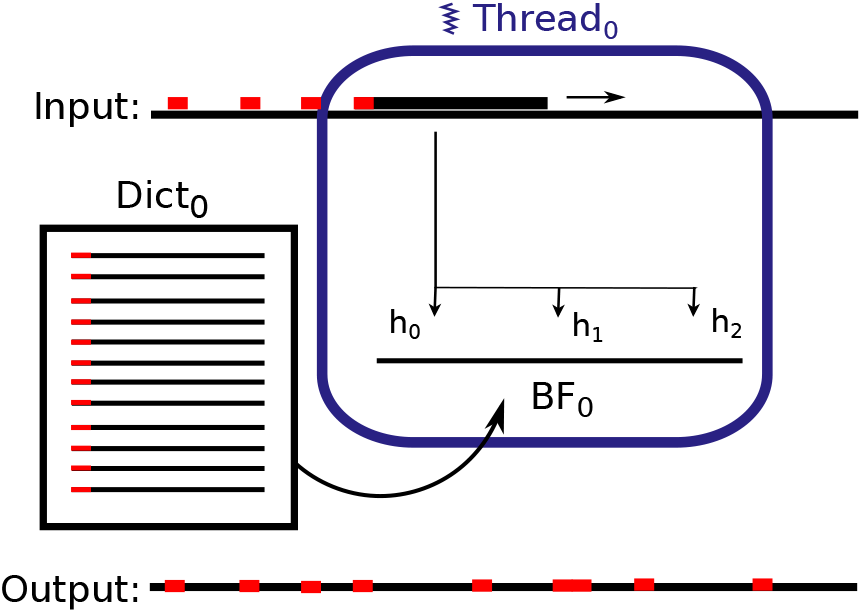
Overview of the long read filter.

Depending on the sensitive position that the filter is able to detect in subsequences, some nucleotides at the end, or at the beginning, of a read may not be studied. These non-studied parts of a read can preventively be treated as sensitive. This is a small price to pay compared to the decreased number of false positives that this technique brings compared to other filtering approaches. Moreover, this effect tends to disappear once we start using several Bloom filters, initialized with different sensitive positions.

#### D. Tolerating sequencing errors

The filtering approach we described earlier in this paper detects all genomic variations embedded in error-free reads. Although sequencing technologies are under constant improvement, it is nowadays still important to tolerate the sequencing errors they may introduce in reads (i.e., insertions, deletions or mutations). Consequently, we now extend the Long Read Filtering (LRF) approach with mechanisms to tolerate sequencing errors.

More precisely we study how we can iterate the error-free detection mechanism to tolerate some sequencing errors. We therefore use several Bloom filters, initialized with different sets of (*K, i*)-sensitive sequences. Intuitively, iterating the detection process increases the probability for a sensitive nucleotide to be detected in presence of errors.

In our implementation, all Bloom filters are queried in parallel by several threads to maximize the overall throughput, as they can all fit into the main memory of the machine used for experiments. Each thread tags a subset of a read’s nucleotides as sensitive, and after all Bloom filters finishes, the union of tagged nucleotides is computed to obtain the final set of nucleotides detected sensitive. Depending on the computing resources one has, it is possible to apply the various filters differently. The spectrum ranges from several machines applying the filters in parallel, for maximal throughput, to one machine applying the filters sequentially, for the lowest hardware requirements.

At initialization time, to explain which sequences should be inserted in the Bloom filters, and which positions in these sequences are considered sensitive, let us consider a sequence of 2*K* − 1 nucleotides, which represents a general situation where one sensitive nucleotide is surrounded by two flankings of *K* − 1 nucleotides. We aim at defining evenly distributed sensitive positions in the interval 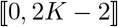.

Let us consider the situation where we define *B* Bloom filters over this set of positions, initialized with sequences of *K* nucleotides. The sensitive position in each of these Bloom filters is the position of the sensitive nucleotide in the subsequence they are built on. If *B* equals 1, any sequence containing the sensitive nucleotide will bring similar results. In this case, we create all sequences where the last nucleotide is sensitive. If *B* is greater than 1, we define 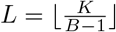 to simplify notations. We define the *B* intervals of nucleotides to be inserted in these Bloom filters as follows:

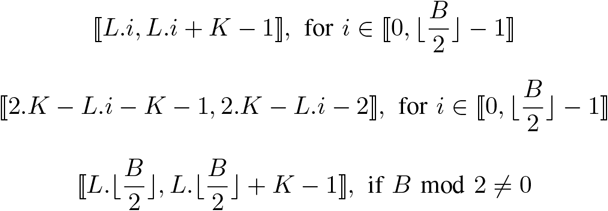

Figure 3 shows an example where, at some point of the filtering process, to detect one sensitive nucleotide, three subsequences of 11 nucleotides (i.e., K=11) are selected and given as input to different Bloom filters. These Bloom filters detect sequences where respectively the last, the first, or the 6th nucleotide is sensitive. In this example, two nucleotides have been altered by errors, a G on the left and another C on the right. The first and second Bloom filters do not detect the corresponding subsequences as sensitive due to the errors. However, the third one does detect it, because there are no errors modifying the nucleotides in the subsequence that this filter studies.

**Fig. 3.**
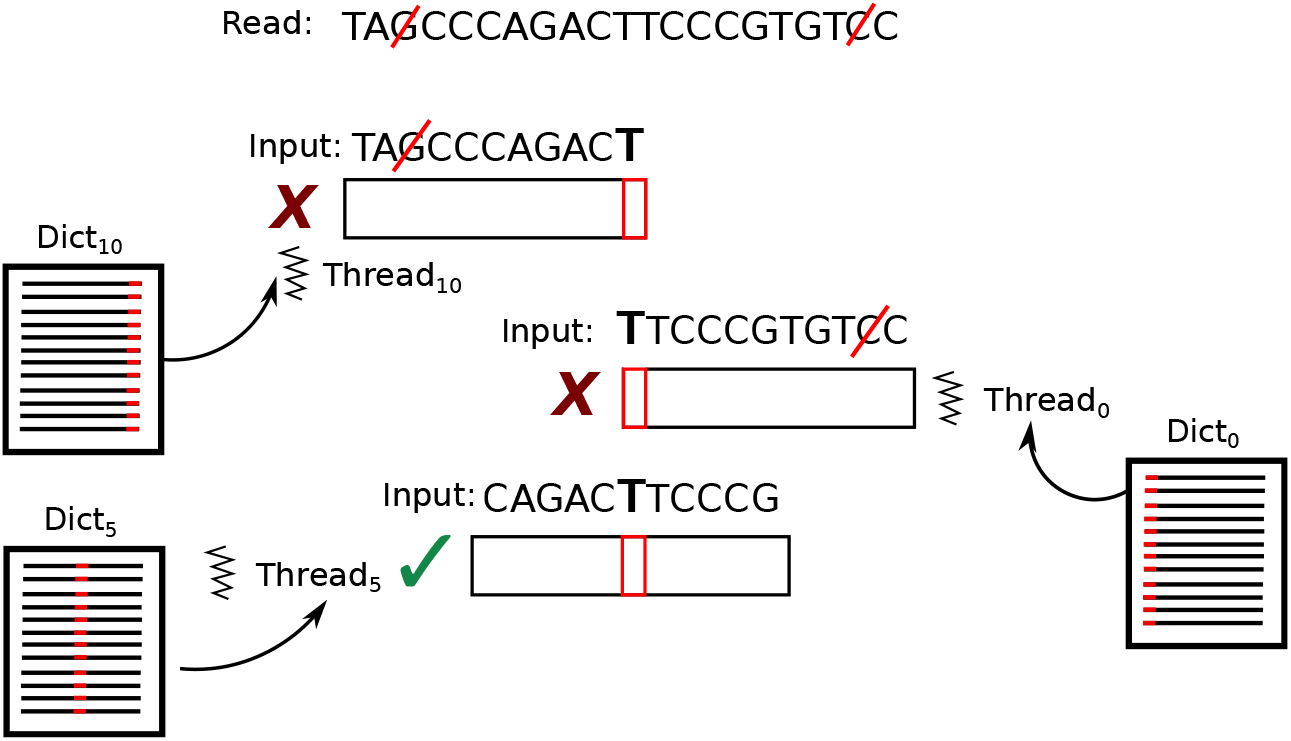
Example of sequences of 11 bases studied by 3 Bloom filters where respectively the last, the first, and the middle position are sensitive. Here, two bases are modified, and only the third sequence would match in the associated dictionary made of 11-bases long sequences whose middle base is sensitive.

## III. RESULTS

In this section, we first describe the system configuration we used for our experiments, using the data sources we presented in Section II-A. Then, we compare the long read filter (LRF), configured with one Bloom filter for fairness, to the short read filter (SRF), which is the only previous filtering approach. We evaluated their performance based on the following criteria:

**Proportion of a genome, or a read, detected sensitive:** The lower this proportion the more information will remain in the reads after filtering of sensitive information;

**False positive rate:** proportion of nucleotides that are detected sensitive and that, in reality, are not sensitive;

**Tolerance to errors:** proportion of sensitive nucleotides detected sensitive in presence of sequencing errors;

**Memory usage:** memory space required to store the files containing the sensitive database, and the size of the corresponding Bloom Filter;

**Throughput:** number of nucleotides that are classified per time unit.

### A. System Configuration

#### Code

We wrote the code that generates the dictionaries of sensitive sequences from the 1000 Genomes database of genomic variations in Python. The LRF and our SRF implementation have been written in C++ for performance. This code executes the following phases: (i) reading of the input file, which contains the raw reads; (ii) initialization of the Bloom filter(s) with the dictionaries of sensitive sequences; and (iii) filtering of the reads.

#### Dictionaries and Bloom filters

We parameterize each Bloom filter with a false positive rate of 10^−3^. This parameter has an impact on the LRF’s false positive rate. However, the LRF’s false positive rate is mainly due to the dictionaries being exhaustive: a sequence may be detected sensitive in an non-sensitive location, because it is sensitive in another genome, or in another location of the genome. The number of elements to be inserted in the Bloom filters is directly given by the dictionaries of sensitive sequences we build. Given these two parameters, the number of bits and the number of hash functions in the Bloom filters are derived using the usual Bloom filter formulas. We choose a value of 10^−3^ because using a lower false positive for Bloom filters does not significantly improve the LRF’s accuracy, and increases its memory consumption. We use MurmurHash [59] to hash sequences for the Bloom filters.

All experiments described in this paper are performed on a quad socket Intel Xeon E5-4650 v3 processor with 12 cores machine running at 2.10 GHz. This machine is equipped with 190 GB RAM, and 15 TB disk storage, which allowed parallel experiments and the generation of all sensitive databases corresponding to different parameters of the filters. One should however note that even with such a powerful machine it was not possible to generate the intermediary files containing the sequences generated with more than 8 combinations of genomic variations.

### B. Proportion of a genome detected sensitive

Figures 4 and 5 show the proportion of nucleotides that are detected sensitive (either true positives or false positives) using one of the two databases built from either genomic variations only or STRs only. For each database, we tested the extended SRF and our approach with one (LRF-1), two (LRF-2) or three (LRF-3) Bloom filters. We explain in the next section which positions these filters study. These two figures also represent the minimum theoretical percentage that can be reached, which is the proportion of nucleotides involved in genomic variations (around 3%) or STRs (around 0.5%). Our approach has the most significant impact on genomic variations where extended SRF cannot classify less than 50% of the nucleotides as sensitive, whereas our approach always converges towards the minimum value. Among the LRF approaches, LRF-1 presents the nearest percentage of nucleotides detected sensitive, which means a small number of false positives. However, LRF-2 and LRF-3 combine both the smaller false positive (i.e., around 10% of a genome is detected sensitive using K=34) and the smaller false negative rates (i.e., less than 10 sensitive nucleotides are not detected per human genome).

**Fig. 4.**
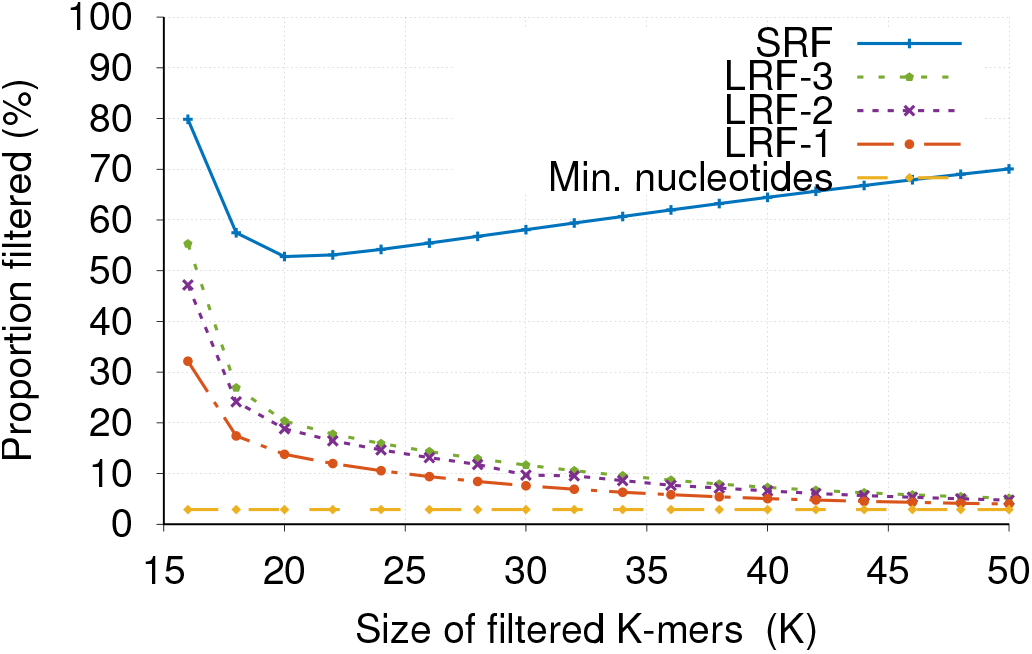
Proportion of nucleotides detected sensitive per K-mer size in a human genome with a filter initialized with the GVs only.

**Fig. 5.**
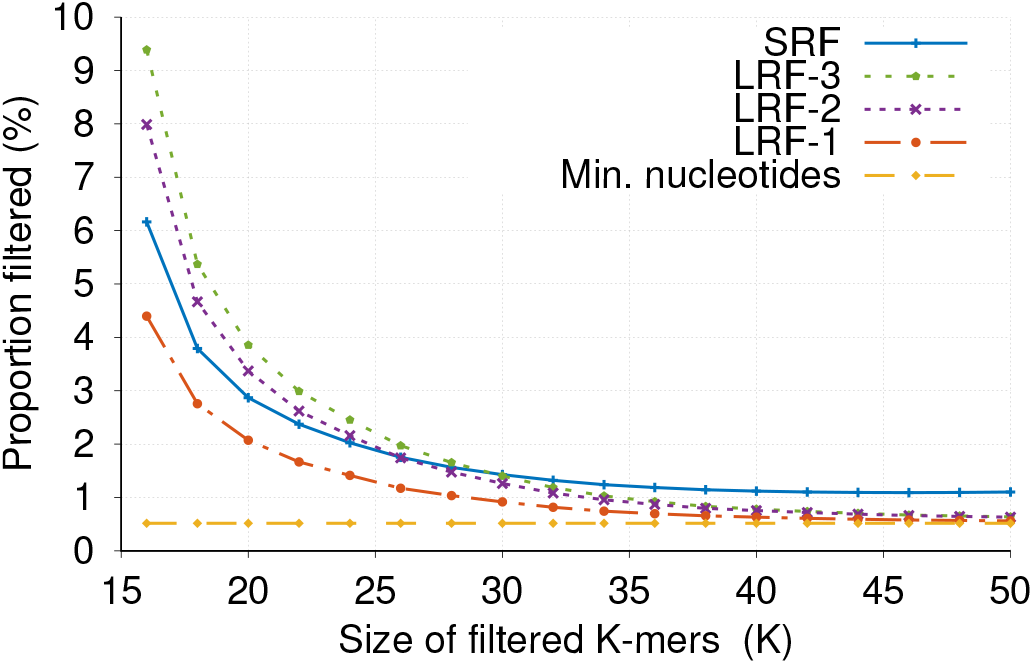
Proportion of nucleotides detected sensitive per K-mer size in a human genome with a filter initialized with the STRs only.

### C. Proportion of reads detected sensitive, or preventively obfuscated

As said in Section II-C, depending on the sensitive position that the filter is able to detect in subsequences, some nucleotides at the end, or at the beginning, of a read may not be studied. These non-studied parts of a read are preventively treated as sensitive. Figure 7 presents the average proportion of reads that is considered sensitive depending on their length. We considered reads of 150, 350, 1000 and 10000 nucleotides, and present the results for the LRF-3 approach. The proportion of reads that is wrongly detected sensitive using LRF is always smaller than the proportion obtained with SRF, which does not have to preventively consider sensitive the extremities of reads. Figure 7 indicate that with short reads (i.e., less than 150 bases), one should initialize its dictionaries with shorter sequences. With reads of 150 bases, the best length for dictionary entries is 20 bases, and the LRF detects 47% of these reads as sensitive. With longer reads, the proportion of reads that is filtered always decreases with the length of the dictionary entries, and we therefore recommend using a length of 34 bases, as justified previously. The longer the reads the smaller the proportion of reads that is detected sensitive, until it reaches the value we obtained with full genomes, which indicates that the LRF approach is more practical with longer reads.

### D. False positive rate

Figure 6 illustrates the percentage of false positive (FP) sensitive nucleotides depending on the size of the subsequences studied. To minimize the number of sensitive nucleotides not detected by the filters, we increased the size of the dictionaries of sensitive sequences (by generating all combinations of genomic variations). Using these larger dictionaries increased the number of false positives, as more sequences in a human genome can be detected sensitive. Experimentally, studying short sequences leads to high FP percentages, between 90% and 100%, for all four approaches. Studying larger sequences still produces a very high FP rate for the SRF, while the FP rates of LRF-1, LRF-2, and LRF-3, respectively, go down to 30%, 40% and 43%. These results show that selecting the largest K value is the best choice to minimize the FP rate. However, with larger sequences the probability to miss a sensitive nucleotide increases, as shown in Appendix, in Figure 12. Decreasing the false positive rate is important because it means that more genomic data can be used to successfully align reads. However, later in the genomic workflow, after reads have been aligned to a reference genome, these false positives would be identified, and removed from the set of sensitive nucleotides.

**Fig. 6.**
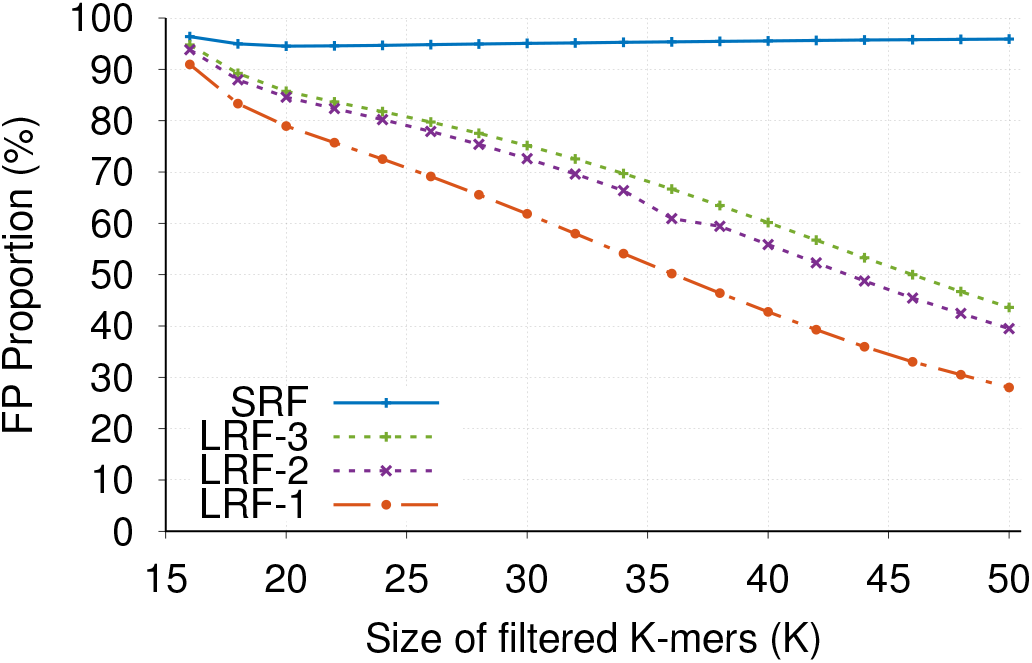
Proportion of false positives on a full genome using SRF [15] and LRF with one (LRF-1), two (LRF-2), or three (LRF-3) Bloom filters, per K-mer size.

**Fig. 7.**
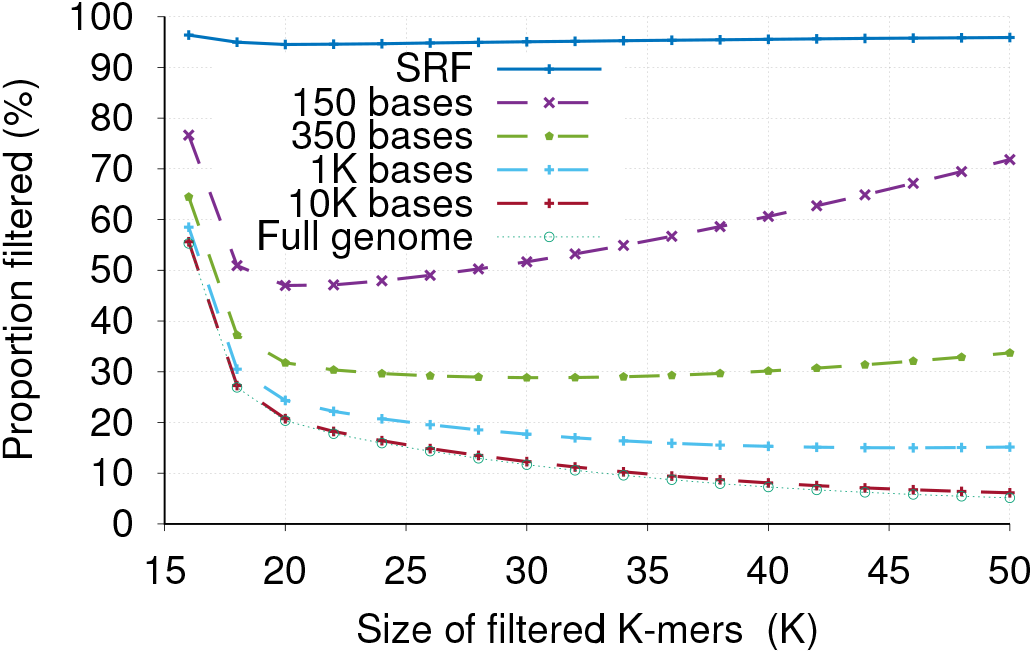
Proportion of nucleotides detected sensitive per read depending on the read length, using LRF-3, including the nucleotides preventively considered sensitive at the extremities of the reads.

### E. Tolerating sequencing errors

Let us consider again a probability of mutation per nucleotide *p*_*m*_. The overall probability *P*_*detect*_ that a sensitive nucleotide is identified by at least one of the *B* Bloom filters can be expressed in function of the probability that the sequence contains M mutations *P*_*mut*_ and the probability that the sensitive nucleotide is detected given M mutations *P*_*detect*|*mut*_ as follows:

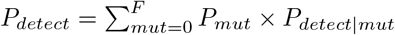

The probability that exactly *M* mutations happen among 2*F* nucleotides follows a binomial distribution, and is equal to 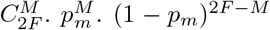. As both *P*_*mut*_ and *P*_*detect*|*mut*_ quickly decrease when *M* increases, it is convenient to evaluate *P*_*detect*_ through simulations. We perform several experiments to determine the probability that a sensitive nucleotide will be correctly detected by a given number of Bloom filters depending on a mutation rate per nucleotide.

Figure 8 presents this detection probability with a 2% error rate, depending on the sizes of the subsequences and the number of Bloom filters used. The rationale behind studying a 2% error rate is that second generation sequencing machines have lower error rates. We believe that the error rate in longer reads will reach or be lower than 2% in the near future. Generally, the shorter the subsequences are the higher is the probability that the sensitive letter will be detected. With sequences of 20, 24, 30 and 34 nucleotides, the probability of detection tends to 0.94, 0.92, 0.88 and 0.84, respectively. However, short subsequences would also increase the proportion of false positives, as we discussed previously. One can also see that when using more than three Bloom filters, the probability of detection increases slowly, which indicates that using three Bloom filters would be the best choice from a sensitivity versus overhead point of view.

**Fig. 8.**
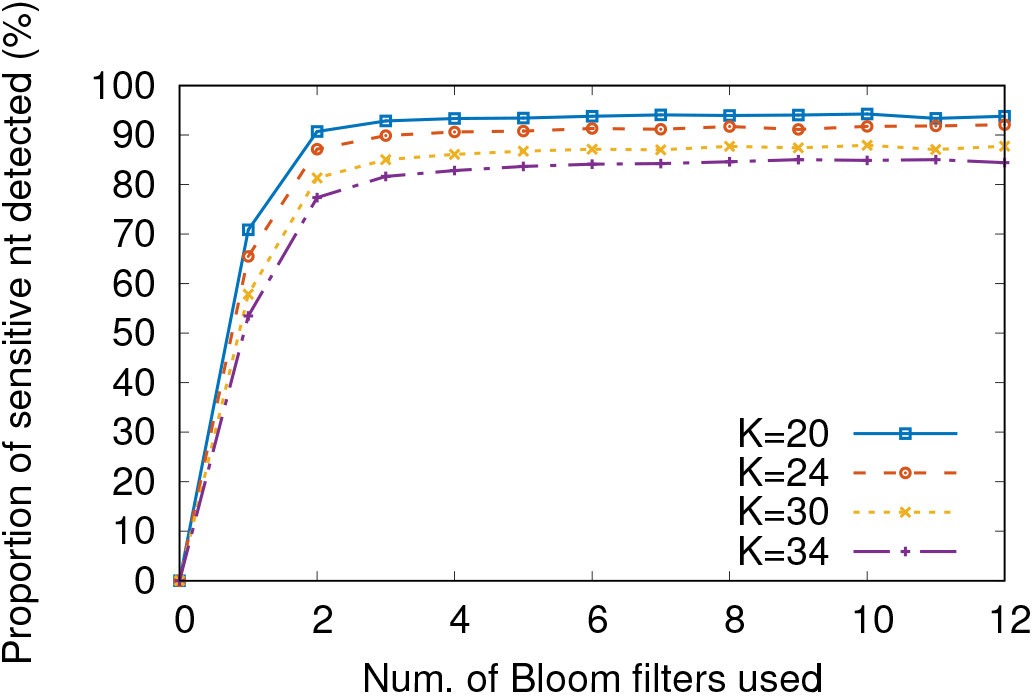
Proportion of sensitive nucleotides detected depending on the size of the K-mers and the number of Bloom filters used with a 2% error rate.

Next, Figure 9 presents the proportion of sensitive nucleotides that are detected depending on several mutation rates, and on the number of Bloom filters used, when considering 30-mers. For this experiment, we used 30-mers, instead of the optimal set of 34-mers, to have a fair comparison against SRF. The higher the error rate, the smaller the proportion of sensitive nucleotides that are detected. The probability for a sensitive nucleotide to be detected when it presents an error rate of 0.1%, 1%, 2% or 4% is 99.97%, 96%, 86% and 62%, respectively. These numbers have to be compared to those of SRF (Section A). For example, with 2% of errors, the LRF approaches detect up to 86% of the sensitive nucleotides while SRF detects 56%. For comparison, with 2% of errors, the LRF approaches detect up to 86% of the sensitive nucleotides while SRF detects 56%.

**Fig. 9.**
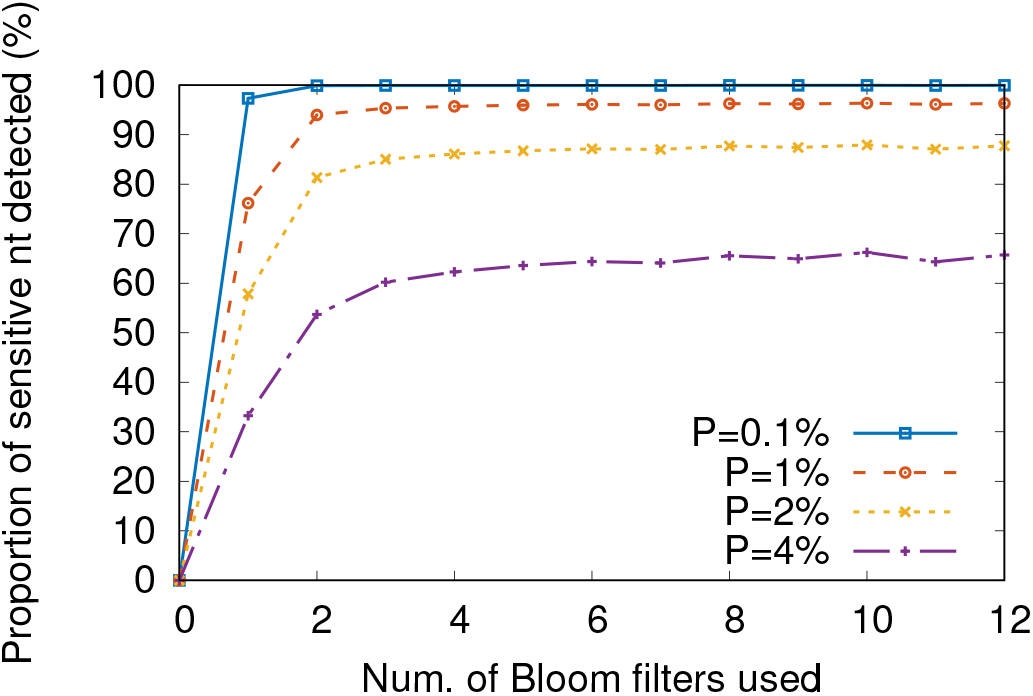
Proportion of sensitive nucleotides detected depending on the error rate and the number of Bloom filters used for 30-mers (SRF’s settings)

### F. Memory Usage

Figure 10 illustrates the size of a Bloom filter in the LRF and the SRF approaches depending on the length of the sensitive sequences that are inserted in the dictionaries.

**Fig. 10.**
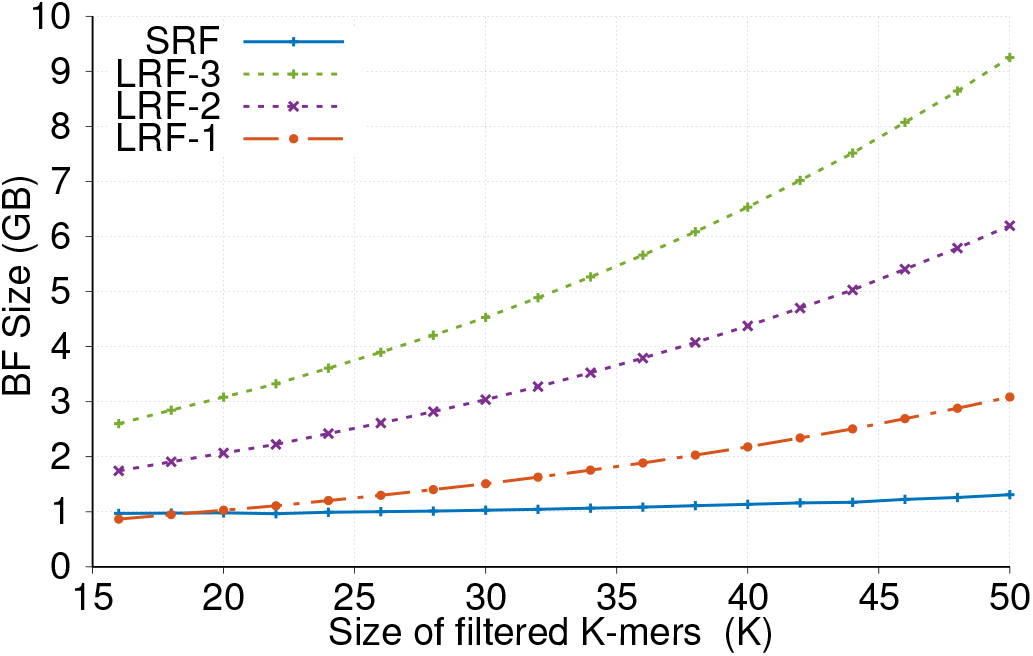
Bloom filter size depending on the length of K-mers with all filtering approaches.

One can observe that the sizes of the Bloom filters increase with the length of the sensitive sequences they are initialized with, which means that more sensitive sequences are generated. It first comes from the fact that a larger proportion of longer sequences contain at least one sensitive base. Second, for the LRF approaches, since longer sequences are more susceptible to contain more genomic variations than shorter ones, there are more combinations of genomic variations. The size of one Bloom filter goes up to 3GB in the LRF-1 approach with 50-mers, while it measures less than 1.5GB with the SRF approach. This difference is due to the combinations of variations that we generate with the LRF approach.

As expected, iterating the LRF approach increases the memory overhead linearly with the number of Bloom filters used. Using the LRF-3 approaches, which uses 3 Bloom filters, require less than 10GB of memory, which is reasonable given modern hardware.

### G. Throughput

To evaluate the throughput of the filtering approaches, we evaluated the filtering methods (LRF and SRF) with sensitive sequences of 34 nucleotides (i.e., *K* = 34). 34 is the smallest value for which the LRF-3 filter produces less than 10% of false positives. Choosing the smallest possible size for the sequences minimizes the size of the Bloom filters, and therefore maximizes the throughput, mainly because less hash functions are needed with smaller Bloom filters. For fairness, we measure the throughput of each approach using only one thread. In practice, one could rely on parallelization to increase the filtering throughput of each approach.

Figure 11 shows the throughput according to different FP rates. As expected, the throughput is higher when the tolerated rate of false positives increases, as the number of hash functions used and the bit table size in Bloom filters decrease. Our approach, LRF-1, has a lower throughput than SRF, which is due to the ways subsequences are studied. Indeed, SRF uses non-overlapping windows in reads, while LRF adopts a sliding window approach. The tradeoff is that the SRF approach is faster, with a throughput ranging from 9.1 to 12 millions of nucleotides per second, while LRF is significantly better with relation to false positives and false negatives. Considering that current sequencing machines have a throughput of 0.3 millions of bp/sec [60], LRF is still ten times faster, with a throughput ranging from 3.3 to 4.9 millions of nucleotides per second, depending on the false positive rates.

**Fig. 11.**
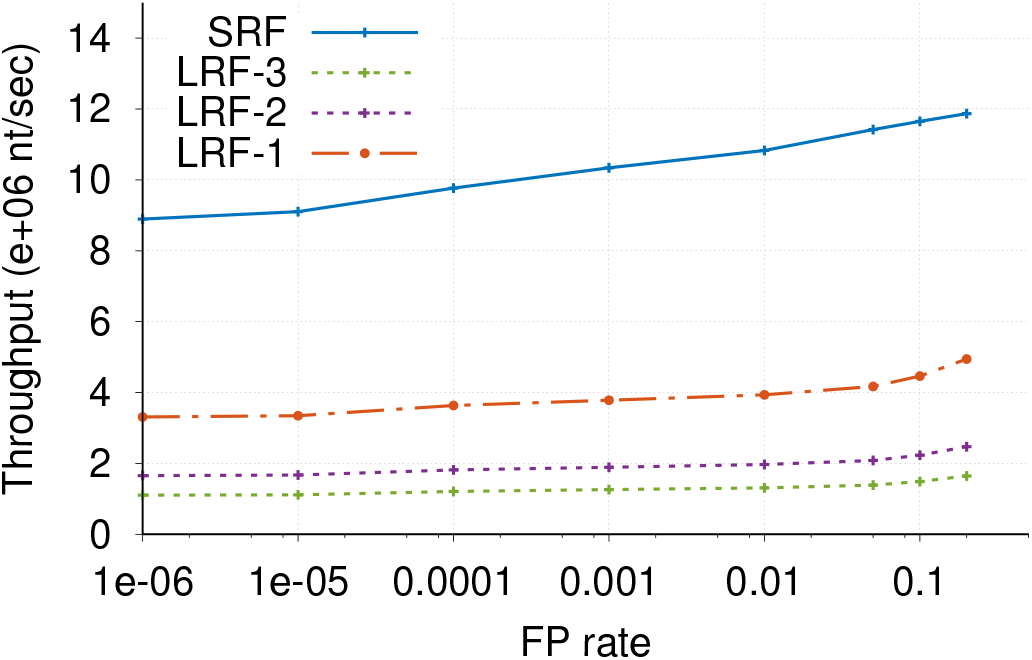
Throughput of the filtering approaches for different false positive rate values with sequences of 34 nucleotides, using one thread.

**Fig. 12.**
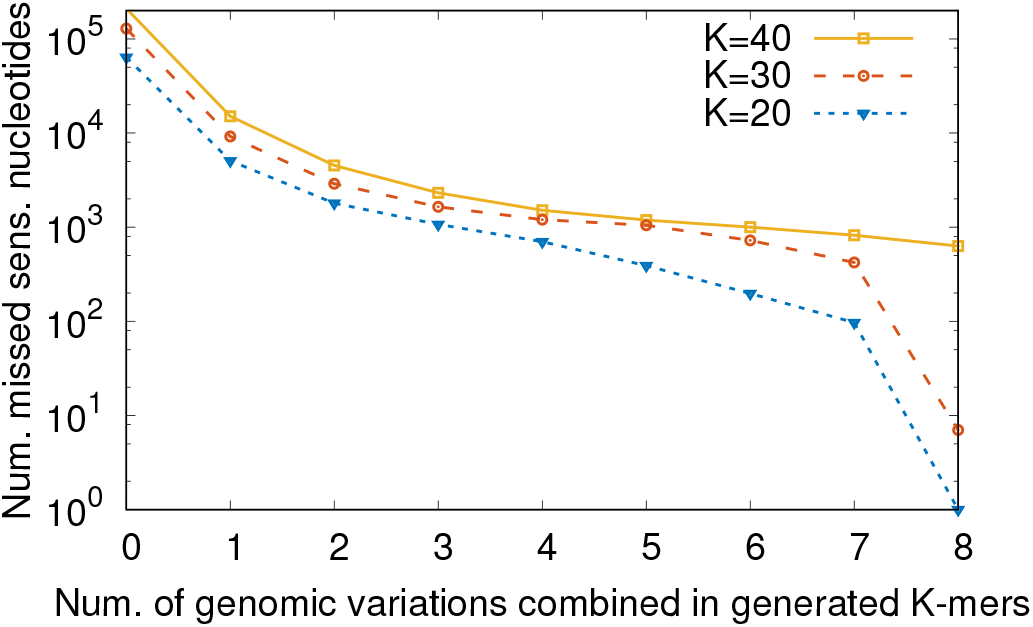
Number of undetected sensitive nucleotides depending on the number of GVs combined when creating sequences of 30 nucleotides.

## IV. DISCUSSION

In this section, we review the limitations of our filtering approach, present the choices we made concerning our methods and experiments, and detail potential impacts of this work on the associated research area.

### A. Limitations of the filtering approach

#### Completeness of genomic variations databases

With high probability, genomic variations that have not yet been identified by the scientific community will not be detected by the filter. However, updating the filter to include new sensitive genomic variations as they are discovered is straightforward: one would have to generate the possible genomic sequences that carry the new genomic variation, and insert them in the existing Bloom filter, which merely consists of hashing the sequences and setting adequate bits to 1. If the number of new sequences to insert becomes too large, it may become necessary to create and initialize a larger filter, since the false positive rate of a Bloom filter increases with the number of elements it contains (we refer to Section I-A6 for more details on Bloom filters). In addition, new genomic variations are now more rarely discovered [15], which limits the residual risk of not detecting a large number of yet unknown sensitive sequences. In case a genomic variation is no longer considered valid removing it from the Bloom filters is more complicated. One would have to reinitialize a new Bloom filter without it, or could rely on counting Bloom filters.

#### Bloom filter detection limit

Per human genome, the filter did not detect less than 10 sensitive nucleotides out of nearly 90 millions. These nucleotides could however be detected using a higher threshold of combined variations locally. Consequently, these few highly variable regions could be treated as special cases to allow for the detection of the genomic variations they contain.

#### Sequencing errors

Tolerating sequencing errors when relying on Bloom filters, which are based on exact matching through hashing, is challenging. In this manuscript, we detail how to use several Bloom filters in parallel to increase the detection accuracy. Currently, the number of Bloom filters used for detection is static. This number of Bloom filter could be dynamically decided depending on the read quality that fastq files contain. Other methods may exist that would improve the sensitivity of the filter in presence of errors. Making use of the whole information contained in reads, through the use of Locality-Sensitive Hashing, and using spaced-Kmers are a possibility we plan to explore in future works.

### B. Decisions made on the methods and experiments

#### Optimal length of K-mers for filtering

We limit the number of genomic variations combined in dictionary entries to 8. We justify in A that it enables both accurate detections and a lower dictionary size. We then found out that the length of the Kmers studied by the Bloom filters has to be lower than 40, otherwise too many sensitive nucleotides are not detected, as supported by Figure 12. However, we also found out that the longer the studied K-mers are the lower the proportion of false positives created by the filter. Therefore, selecting K-mers of 34 nucleotides is an adequate choice to create and detect sensitive nucleotides.

#### Reducing the size of Bloom filters

In theory, it would be possible to reduce the number of sequences produced if we avoid creating sequences that contain a configuration of alleles that is very unlikely. Using linkage disequilibrium (LD) to discard improbable sequences reduces the number of generated sequences per dictionary from 2 to 5% depending on the length of the sequences (the longer the sequences the more are discarded). Relying on generalized linkage disequilibrium metrics would probably further optimize this result.

### C. Potential impacts of this work on the area

We foresee that previous works related to privacy-preserving read alignment will benefit from the use of a long read filter, such as the one we presented in this manuscript.

Currently, due to the rapid advances of sequencing technologies, biocenters tend to use Clouds to store and align reads [61]. Plaintext approaches, like BLAST [62] have been parallelized (e.g., CloudBLAST [63]) to improve their performance, when numerous CPUs or machines are available. However, a user relying on these algorithms to align sensitive genomic data needs to trust the cloud provider, which does not provide security guarantees [19]. Given the long duration for which genomic data remains sensitive, this solution is not satisfactory from a privacy point of view. In contrast to the plaintext alignment procedures, our filter is lightweight, fast, and can be run directly on a sequencing machine without being a bottleneck in the genomic pipeline.

Our filter would limit the use of such algorithms to data that is detected sensitive, and would improve the overall performance of a privacy-preserving alignment. Currently, we envision that our filter approach could be used to obfuscate sensitive data from reads, releasing only the reads that can be safely aligned in public clouds in plaintext, while the reads that could not be aligned in public clouds would have to be aligned a second time, using their non-obfuscated versions, either on a private cloud, or using secure alignment algorithms. The precise design and performance evaluation of this envisioned alignment scheme, which our filter would enable, is future work.

## V. CONCLUSION

In this paper, we presented a novel filtering approach to detect sensitive nucleotides in long reads, which are now produced by the newest sequencing technologies. Our methods improve on the state of the art both in terms of false negatives (from 100,000 to less than 10 missed sensitive nucleotides) and false positives (10% of a genome is filtered instead of 60%). We also studied how well this method performs in presence of sequencing errors, and showed that iterating the detection process helps tolerating such errors (86% of the sensitive nucleotides are detected instead of 56%, with 2% of errors). In addition, despite its non-negligible memory and CPU overhead, the filtering process is still faster than current sequencing machines, and can be parallelized in several ways, suggesting that our method will easily scale-up with the throughput evolution of the former. This privacy filter produces reads where sensitive information has been masked, therefore protecting the privacy of subjects in the standard genomic workflows, where reads are often aligned to a reference genome. Our filter can be applied to reads of any size and is designed to detect all kinds of genomic variations, missing less than 10 sensitive nucleotides per genome.

Common to all automated early filtering approaches, our method allows: (i) the incorporation or merging of filtering in a straightforward way to NGS machines architectures with local (protected) storage; (ii) the storage of sensitive information per subject in a separate and more secure location, to be added back to reads only when desired or needed, e.g., after the alignment step. Future work includes the in-depth study of masked reads alignment to further improve this result.

## ACKNOWLEDGEMENTS

This work was supported by the Fonds National de la Recherche Luxembourg (FNR) through PEARL grant FNR/P14/8149128, and by the Fundação para a Ciência e para a Tecnologia (FCT) through funding of the LaSIGE Research Unit, ref. UID/CEC/00408/2013.

## APPENDIX

As Figure 13 shows, the flanking sequences of variations often contain variations themselves. To generate this figure, we studied all genomic variations of the 1000 genomes project and counted their neighboring variations located less than 30 nucleotides away. We then plotted the cumulative distribution of this number of close-by variations. From those results, 60% of the genomic variations are surrounded by at least one other genomic variation and 99.96% of the sequences contain less than 8 genomic variations (on both sides). Therefore, a sensitive genomic variation or an STR of a specific subject may not be detected by the SRF, if its flankings contain at least one genomic variation that is not shared with the reference genome.

**Fig. 13.**
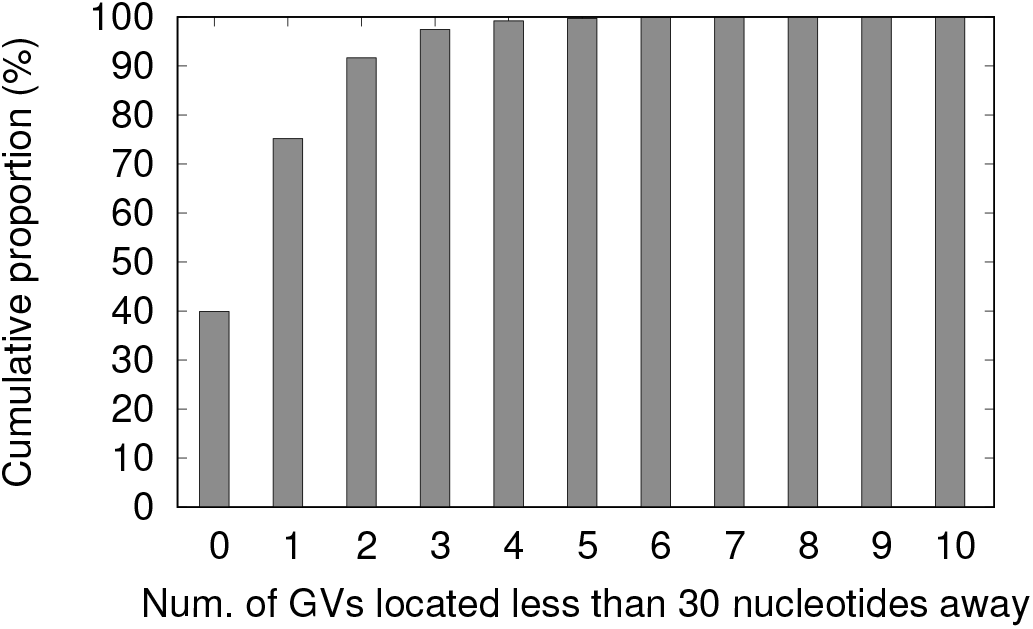
Cumulative distribution of the number of genomic variations contained in sensitive sequences of 30 nucleotides.

Figure 12 justifies our choice of combining up to 8 genomic variations. This figure presents the average number of sensitive nucleotides that are not detected in a non-reference genome (using a logarithmic scale) depending on the number of variations combined during the generation of the dictionary. We studied three different lengths of subsequences to present to our filter (20, 30 and 40). For lengths of 20 or 30 nucleotides and with 8 genomic variations involved in combinations, less than 10 sensitive nucleotides are missed per genome, while a filter that would not consider combinations would miss around 100,000 nucleotides. Using longer, 40-nucleotide sequences deteriorates detection performance, making the filter miss around 600 variations, as longer sequences are more susceptible to contain more variations.

In this appendix, we explain why the short read filter [15], which we call SRF for short, cannot be applied to long reads. Such infeasibility demonstrates the novelty of our contribution, the long read filter, which enables the filtering of long reads, and therefore opens new research directions.

### Infeasibility of combining genomic variations with SRF

Bloom filters are based on exact matching, and therefore their output is very sensitive to incorrect or incomplete initialization. To solve this issue, we combine the alleles of several genomic variations when computing the sensitive sequences. It is important to note that we would not have been able to generate such combinations of variations with the SRF approach, because it would consume too much disk space (more than 20TB) and/or would probably be very slow. The reason behind this is that the SRF dictionary basically contains all the possible dictionaries that our long read filter could build.

### Infeasibility of obtaining an acceptable false positive rate with SRF and long reads

SRF classifies sequences of 30 nucleotides as sensitive or non-sensitive and cannot be directly extended to longer reads, which are becoming more and more common. Indeed, as Figure 14 shows, the proportion of sequences that contain at least one sensitive nucleotide grows very quickly with their size. For example, 88% of the 100-nucleotides reads and 94% of the 200-nucleotides reads would be considered sensitive. As it is clearly not practical for research purposes to classify nearly all reads as sensitive, or to create a dictionary of all possible sensitive long reads, it is now necessary to filter reads at the level of their nucleotides.

**Fig. 14.**
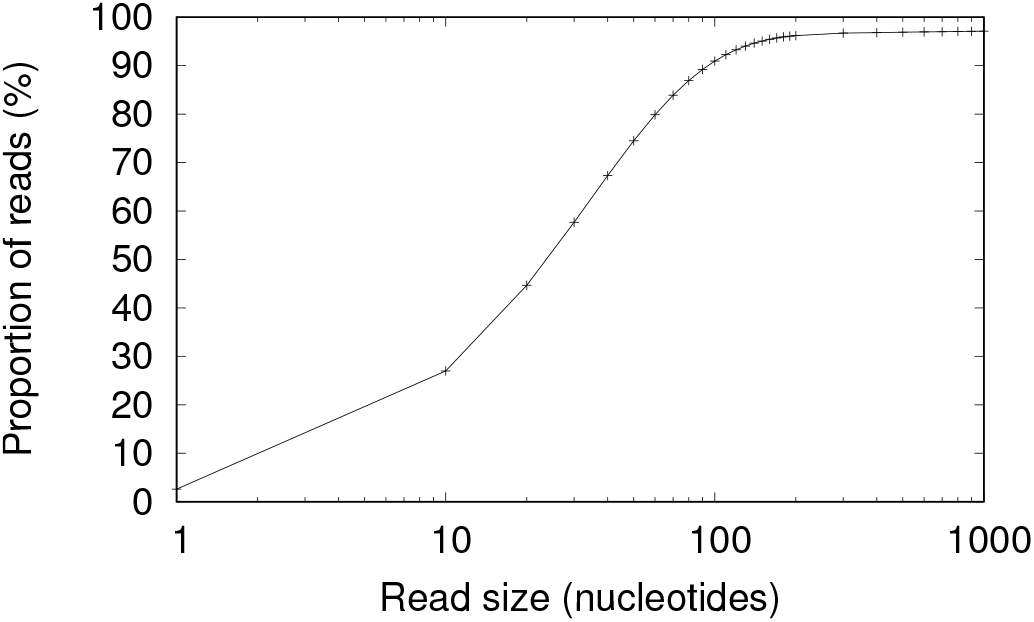
Proportion of reads containing at least one sensitive nucleotide depending on their length (logarithmic scale)

The first idea to extend SRF to long reads consists in studying each non-overlapping subsequence of a long read. In such a scheme, a sensitive nucleotide (in bold in the top line) would be included in exactly one studied subsequence, which therefore would be considered sensitive, along with all the other nucleotides in the subsequence. This approach would detect sensitive nucleotides since sensitive sequences built from the genomic variations or STRs include the nucleotides that precede and follow sensitive portions of the genome. This idea is the one Cogo et al. [15] implicitly used to evaluate the proportion of sensitive nucleotides in several human genotypes. We therefore consider this as the baseline approach, and will refer to it as SRF, when discussing and evaluating the extension to longer reads. However, the granularity of this approach is not perfect: if a window is declared sensitive by the filter then all its nucleotides also have to be considered sensitive, because the position of the sensitive nucleotide in the subsequence is unknown. In practice, this method generates many false positives, as the flankings of a genomic variation or of an STR, which are needed to detect the variations, are declared sensitive.

The short read filter could also be applied in various sliding window fashions (i.e., studying overlapping subsequences), but that would, however, create even more false positives or produce random results in case of sequencing errors.

### Sequencing errors and sensitivity

Even though sequencing technologies are under constant improvement, sequencing machines can introduce mutations and insertion/deletions in the reads they produce. Due to the exact matching property of Bloom filters, the short read filtering approach is not able to detect sensitive nucleotides that are located near a sequencing error. However, these nucleotides are still sensitive, so aligning a read, by e.g., using one of the alignment algorithms tolerating such sequencing errors, such as BWA [2] or Bowtie [64], would allow an attacker to obtain the genomic variations.

Applying SRF, a sensitive character is detected only if the 30-nucleotides subsequence it belongs to is detected sensitive. If any nucleotide of this window is modified, or deleted, then there are high chances that the sequence and thus the sensitive nucleotide will no longer be detected sensitive. Indeed, let *p*_*m*_ be the probability that a nucleotide is modified by a sequencing machine. Then, the probability that all 29 nucleotides inside a SRF window are not modified is equal to (1 − *p*_*m*_)^29^. With 0.1%, 1%, 2%, or 4% of mutation errors, the SRF would then detect respectively, 97%, 75%, 56%, and 31% of the sensitive nucleotides, possibly making thousands of genomic variations available for an attack.

## REFERENCES

[1] S. E. Levy, R. M. Myers, Advancements in next-generation sequencing, Annual Review of Genomics and Human Genetics 17 (1) (2016) 95–115.

[2] H. Li, R. Durbin, Fast and accurate short read alignment with burrows–wheeler transform, Bioinformatics 25 (14) (2009) 1754–1760.

[3] B. Langmead, C. Trapnell, M. Pop, S. L. Salzberg, et al., Ultrafast and memory-efficient alignment of short dna sequences to the human genome, Genome biol 10 (3) (2009) R25.

[4] R. Wang, Y. F. Li, X. Wang, H. e. a. Tang, Learning your identity and disease from research papers: information leaks in genome wide association study, in: ACM CCS, 2009.

[5] A. J. Pakstis, W. C. Speed, R. Fang, F. C. e. a. Hyland, Snps for a universal individual identification panel, Human genetics 127 (3) (2010) 315–324.

[6] K. Kidd, A. Pakstis, W. Speed, E. e. a. Grigorenko, Developing a snp panel for forensic identification of individuals, Forensic science international 164 (1) (2006) 20–32.

[7] M. Gymrek, A. L. McGuire, D. Golan, E. e. a. Halperin, Identifying personal genomes by surname inference, Science 339 (6117) (2013) 321–324.

[8] M. Humbert, K. Huguenin, J. Hugonot, E. Ayday, J.-P. Hubaux, De-anonymizing genomic databases using phenotypic traits, PoPETS (2) (2015) 99–114.

[9] D. R. Nyholt, C.-E. Yu, P. M. Visscher, On jim watson’s apoe status: genetic information is hard to hide, European Journal of Human Genetics 17 (2009) 147–149.

[10] N. Homer, S. Szelinger, M. Redman, D. e. a. Duggan, Resolving individuals contributing trace amounts of dna to highly complex mixtures using high-density snp genotyping microarrays, PLoS Genet 4 (8) (2008) e1000167.

[11] S. Shringarpure, C. Bustamante, Privacy risks from genomic data-sharing beacons, The American Journal of Human Genetics 97 (5) (2015) 631–646.

[12] M. Backes, P. Berrang, M. Humbert, P. Manoharan, Membership privacy in microrna-based studies, in: ACM CCS, 2016.

[13] E. Ayday, J. L. Raisaro, U. Hengartner, A. Molyneaux, J.-P. Hubaux, Privacy-preserving processing of raw genomic data, in: DPM, 2014, pp. 133–147.

[14] K. Zhang, X. Zhou, Y. Chen, X. Wang, Y. Ruan, Sedic: privacy-aware data intensive computing on hybrid clouds, in: Proceedings of the 18th ACM conference on Computer and communications security, ACM, 2011, pp. 515–526.

[15] V. V. Cogo, A. Bessani, F. M. Couto, P. Verissimo, A high-throughput method to detect privacy-sensitive human genomic data, in: Proceedings of the 14th ACM Workshop on Privacy in the Electronic Society, ACM, 2015, pp. 101–110.

[16] M. J. Atallah, F. Kerschbaum, W. Du, Secure and private sequence comparisons, in: Proceedings of the 2003 ACM workshop on Privacy in the electronic society, ACM, 2003, pp. 39–44.

[17] J. Baron, K. El Defrawy, K. Minkovich, R. Ostrovsky, E. Tressler, 5pm: Secure pattern matching, in: SCN, Springer, 2012, pp. 222–240.

[18] Y. Huang, D. Evans, J. Katz, L. Malka, Faster secure two-party computation using garbled circuits., in: USENIX Security, 2011.

[19] F. L. Rocha, M. Correia, Lucy in the sky without diamonds: Stealing confidential data in the cloud, 2011 IEEE/IFIP 41st International Conference on Dependable Systems and Networks Workshops (DSN-W) (2011) 129–134.

[20] A. Michalas, N. Paladi, C. Gehrmann, Security aspects of e-health systems migration to the cloud, in: e-Health Networking, Applications and Services (Healthcom), 2014 IEEE 16th International Conference on, IEEE, 2014, pp. 212–218.

[21] B. Fabian, T. Ermakova, P. Junghanns, Collaborative and secure sharing of healthcare data in multi-clouds, Information Systems 48 (2015) 132–150.

[22] Y. Tong, J. Sun, S. S. Chow, P. Li, Cloud-assisted mobile-access of health data with privacy and auditability, IEEE Journal of biomedical and health Informatics 18 (2) (2014) 419–429.

[23] B. Yüksel, A. Küpçü, Ö. Özkasap, Research issues for privacy and security of electronic health services, Future Generation Computer Systems 68 (2017) 1–13.

[24] M. T. Goodrich, The mastermind attack on genomic data, in: Security and Privacy, 2009 30th IEEE Symposium on, IEEE, 2009, pp. 204–218.

[25] L. T. Vaszar, M. K. Cho, T. A. Raffin, Privacy issues in personalized medicine, Pharmacogenomics 4 (2) (2003) 107–112.

[26] R. B. Altman, T. E. Klein, Challenges for biomedical informatics and pharmacogenomics, Annual review of pharmacology and toxicology 42 (1) (2002) 113–133.

[27] M. Naveed, E. Ayday, E. W. Clayton, J. Fellay, C. A. Gunter, J.-P. Hubaux, B. A. Malin, X. Wang, Privacy in the genomic era, ACM Computing Surveys (CSUR) 48 (1) (2015) 6.

[28] X. Jiang, Y. Zhao, X. Wang, B. Malin, S. Wang, L. Ohno-Machado, H. Tang, A community assessment of privacy preserving techniques for human genomes, BMC medical informatics and decision making 14 (Suppl 1) (2014) S1.

[29] B. Malin, L. Sweeney, Determining the identifiability of dna database entries., in: Proceedings of the AMIA Symposium, American Medical Informatics Association, 2000, p. 537.

[30] B. Malin, L. Sweeney, How (not) to protect genomic data privacy in a distributed network: using trail re-identification to evaluate and design anonymity protection systems, Journal of biomedical informatics 37 (3) (2004) 179–192.

[31] Z. Lin, M. Hewett, R. B. Altman, Using binning to maintain confidentiality of medical data., in: Proceedings of the AMIA Symposium, American Medical Informatics Association, 2002, p. 454.

[32] B. Malin, Protecting dna sequence anonymity with generalization lattices, Carnegie Mellon University, School of Computer Science [Institute for Software Research International], 2004.

[33] M. Kantarcioglu, W. Jiang, Y. Liu, B. Malin, A cryptographic approach to securely share and query genomic sequences, IEEE Transactions on information technology in biomedicine 12 (5) (2008) 606–617.

[34] B. A. Malin, An evaluation of the current state of genomic data privacy protection technology and a roadmap for the future, Journal of the American Medical Informatics Association 12 (1) (2005) 28–34.

[35] G. Loukides, A. Gkoulalas-Divanis, B. Malin, Anonymization of electronic medical records for validating genome-wide association studies, Proceedings of the National Academy of Sciences 107 (17) (2010) 7898–7903.

[36] M. Humbert, E. Ayday, J.-P. Hubaux, A. Telenti, Addressing the concerns of the lacks family: quantification of kin genomic privacy, in: Proceedings of the 2013 ACM SIGSAC conference on Computer & communications security, ACM, 2013, pp. 1141–1152.

[37] B. H. Bloom, Space/time trade-offs in hash coding with allowable errors, Commun. ACM 13 (7) (1970) 422–426.

[38] M. Akgün, A. O. Bayrak, B. Ozer, M. Şamil Saăıroălu, Privacy preserving processing of genomic data: A survey, Journal of Biomedical Informatics 56 (2015) 103–111.

[39] K. Zhang, X. Zhou, Y. Chen, X. Wang, Y. Ruan, Sedic: Privacy-aware data intensive computing on hybrid clouds, in: ACM CCS, 2011.

[40] Y. Chen, B. Peng, X. Wang, H. Tang, Large-scale privacy-preserving mapping of human genomic sequences on hybrid clouds., in: NDSS, 2012.

[41] V. Popic, S. Batzoglou, Privacy-preserving read mapping using locality sensitive hashing and secure kmer voting, bioRxiv.

[42] J. H. Cheon, M. Kim, K. Lauter, Homomorphic computation of edit distance, in: Financial Cryptography and Data Security, 2015, pp. 194–212.

[43] Y. Erlich, A. Narayanan, Routes for breaching and protecting genetic privacy, Nature Genetics 15 (6) (2014) 409–421.

[44] D. Greenbaum, A. Sboner, X. J. Mu, M. Gerstein, Genomics and privacy: implications of the new reality of closed data for the field, PLoS Comput Biol 7 (12) (2011) e1002278.

[45] L. Sweeney, k-anonymity: A model for protecting privacy, International Journal of Uncertainty, Fuzziness and Knowledge-Based Systems 10 (05) (2002) 557–570.

[46] G. Li, Y. Wang, X. Su, Improvements on a privacy-protection algorithm for dna sequences with generalization lattices, Computer methods and programs in biomedicine 108 (1) (2012) 1–9.

[47] J. D. Thompson, D. G. Higgins, T. J. Gibson, Clustal w: improving the sensitivity of progressive multiple sequence alignment through sequence weighting, position-specific gap penalties and weight matrix choice, Nucleic acids research 22 (22) (1994) 4673–4680.

[48] Z. Zhang, S. Schwartz, L. Wagner, W. Miller, A greedy algorithm for aligning dna sequences, Journal of Computational biology 7 (1-2) (2000) 203–214.

[49] S. Wan, M.-W. Mak, S.-Y. Kung, Protecting genomic privacy by a sequence-similarity based obfuscation method, arXiv preprint arXiv:1708.02629.

[50] A. Broder, M. Mitzenmacher, Network applications of bloom filters: A survey, Internet mathematics 1 (4) (2004) 485–509.

[51] R. Schnell, T. Bachteler, J. Reiher, Privacy-preserving record linkage using bloom filters, BMC medical informatics and decision making 9 (1) (2009) 41.

[52] E. A. Durham, M. Kantarcioglu, Y. Xue, C. Toth, M. Kuzu, B. Malin, Composite bloom filters for secure record linkage, IEEE transactions on knowledge and data engineering 26 (12) (2014) 2956–2968.

[53] S. M. Randall, A. M. Ferrante, J. H. Boyd, J. K. Bauer, J. B. Semmens, Privacy-preserving record linkage on large real world datasets, Journal of biomedical informatics 50 (2014) 205–212.

[54] M. Kuzu, M. Kantarcioglu, E. Durham, B. Malin, A constraint satisfaction cryptanalysis of bloom filters in private record linkage, in: International Symposium on Privacy Enhancing Technologies Symposium, Springer, 2011, pp. 226–245.

[55] R. Schnell, C. Borgs, Secure privacy preserving record linkage of large databases by modified bloom filter encodings, International Journal for Population Data Science 1 (1).

[56] D. Vatsalan, P. Christen, E. Rahm, Scalable multi-database privacy-preserving record linkage using counting bloom filters, arXiv preprint arXiv:1701.01232.

[57] 1000 genomes project, http://www.internationalgenome.org/.

[58] Tandem repeats database, https://tandem.bu.edu/cgi-bin/trdb/trdb.exe.

[59] A. Appleby, Murmurhash 2.0 (2008).

[60] L. Liu, Y. Li, S. Li, N. Hu, Y. He, R. Pong, D. Lin, L. Lu, M. Law, Comparison of next-generation sequencing systems, BioMed Research International 2012.

[61] B. Langmead, M. C. Schatz, J. Lin, M. Pop, S. L. Salzberg, Searching for snps with cloud computing, Genome Biology 10 (11) (2009) 1–10.

[62] S. F. Altschul, W. Gish, W. Miller, E. W. Myers, D. J. Lipman, Basic local alignment search tool, Journal of molecular biology 215 (3) (1990) 403–410.

[63] A. Matsunaga, M. Tsugawa, J. Fortes, Cloudblast: Combining mapreduce and virtualization on distributed resources for bioinformatics applications, in: IEEE ESCIENCE, 2008.

[64] B. Langmead, S. L. Salzberg, Fast gapped-read alignment with bowtie 2, Nature methods 9 (4) (2012) 357–359.

